# Bee Microbiomes Harbor Diverse Antimicrobial Resistance Genes on Plasmids

**DOI:** 10.1101/2025.09.24.678402

**Authors:** Dino Lorenzo Sbardellati, Rachel Lee Vannette

**Affiliations:** Department of Entomology and Nematology, University of California Davis, Davis, California, USA

**Keywords:** Antibiotics, Antibiotic resistance, Plasmids, Bees

## Abstract

Antimicrobial resistance (AMR) is an emerging public health threat. In North America, tetracycline and macrolide antibiotics are often used to prevent or treat bacterial infections in honey bees. Previous research has shown that this practice has led to widespread drug resistance in honey bee gut microbiomes. However, where bee-associated bacteria encode AMR, genomically or on mobile genetic elements, is less well understood. Moreover, how the abundance, diversity, and mechanism of AMR differs between managed honey bees and other bees remains largely unexplored. Here we use existing metagenomic data from two previous studies to profile the AMR genes associated with managed honey bees, commercially produced bumble bees, and wild bumble bees. Our results suggest that honey bee associated bacteria house a greater diversity of AMR genes, specifically on plasmids, compared to bumble bees. In addition, we show that honey and bumble bee bacteria likely develop resistance to tetracyclines via different mechanisms. Overall, this study showcases how agricultural management has shaped the AMR genes associated with bees, and offers insights into the ecological context of differential AMR evolution within host-associated systems.

## Introduction

Antimicrobial resistance (AMR) is an emerging public health concern. In 2019, antimicrobial resistant bacteria were estimated to be directly responsible for 1.27 million deaths, and to have contributed to 4.95 million deaths, globally^1^. A key factor driving the rise of AMR is the overuse and misuse of antibiotics in humans and agricultural systems^2,3^, which act as linked systems in the One Health framework^4^. AMR can arise from mutations in existing genes or through the transfer of mobile genetic elements, such as phage and plasmids, which encode resistance genes^5,6^. Given the growing threat of AMR, it is crucial to understand how antibiotic usage drives not only what types of AMR bacteria acquire, but also where those AMR genes are located (i.e genom9cally or on transmissible mobile genetic elements).

Livestock, including managed bees, are commonly treated with antibiotics, including tetracyclines and macrolides. Honey bees specifically are intensively managed with antibiotics to prevent or treat brood diseases, such as American foulbrood (AFB) and European foulbrood (EFB). This routine use of antibiotics represents a strong selective pressure promoting the emergence and maintenance of AMR genes in bee-associated microbiomes^7^. Indeed, tetracycline resistant strains of *Melissococcus plutonius* and *Paenabacillus larvae*, the causative agents of EFB and AFB, have been isolated from bees sampled in British Columbia^8^ and numerous US states^9^. While there has been research into the AMR genes associated with honey bees, little research has examined the AMR genes associated with other bee species. Comparing how the abundance, diversity, and location of AMR genes in managed bees compares to unmanaged wild bees will shed light on how antibiotic use and human intervention influence AMR development and transmission in bee microbiomes.

Here we leverage publicly available metagenomes from two previous studies^10,11^ to profile the AMR genes associated with managed honey bees, commercially produced bumble bees, and wild bumble bees. We hypothesize that AMR genes will be most prominent in managed honey bees and commercially produced bumble bees, while wild bumble bee microbiomes will contain the lowest amount of AMR genes. We test this hypothesis by comparing how the diversity and abundance of AMR differs across bee populations. We then delve into which AMR genes are encoded in bacterial genomes and which are found on mobile genetic elements (plasmids and phage).

## Methods

### Metagenome Sampling

Briefly, we used genomic detection of AMR from gut metagenomes acquired from bumble bees and honey bees. Paired-end metagenomic read libraries from Hotchkiss et al.^11^ and Sbardellati and Vannette^10^ were downloaded from NCBI using NCBI datasets^12^. The Hotchkiss et al.^11^ dataset was subset to only wild (n=18) and commercial (n=6) *Bombus impatiens* bumble bees. Meanwhile, the Sbardellati and Vannette^10^ dataset was subset to only total metagenomes from commercial *B. impatiens* (n=9) and managed *Apis mellifera* honey bees (n=9). To the best of our knowledge, none of the wild bumble bees in our final dataset have been treated with antibiotics. In contrast, the managed honey bees were sampled from hives which have likely been treated with both tetracycline and tylosin antibiotics. Whether commercially produced bumble bees are treated with antibiotics during production is unknown, but unlikely given the dearth of bacterial pathogens recognized to infect bumble bees^13^.

### Bioinformatic processing

Raw reads were trimmed and quality filtered using Trim-Galore^14^ and Trimmomatic^15^. Reads aligning to the genomes of *A. mellifera* (GCF_003254395.2) or *B. impatiens* (GCF_000188095.3) were removed using Bowtie2^16^. Cleaned reads were assembled using metaSPADES^17^ and mobile genetic elements (either phage or plasmids) were identified via geNomad^18^. Putative phage sequences predicted to be <50% complete by CheckV^19^ were removed.

BBsplit.sh^20^ was used to sort reads in our original cleaned metagenomic libraries. Reads were considered phage if they mapped to phage, plasmid if they mapped to plasmid, and genomic if they did not map to either phage or plasmids. The presence and abundance of antimicrobial resistance (AMR) genes were then predicted by aligning sorted read libraries against the MEGARes database using the program AMR++^21^. Abundance of SNP-verified hits were normalized to number of read hits per million reads.

### Data visualization and Statistical analysis in R

All statistical analyses and data visualizations were conducted in R v4.2.3^22^. Plots were generated using ggplot2. How the proportion of read libraries composed of plasmids changed in response to bee type, how the diversity of AMR genes differed across bee and DNA type, and how the total abundance of genes conferring resistance to specific antibiotic types (aminoglycosides and MLS) were compared between bees using ANOVA and TukeyHSD tests.

## Results and Discussion

### Plasmids Make Up a Greater Proportion of Honey bee Metagenomes

To examine the overall abundance of mobile genetic elements (MGEs) in our dataset, we compared the proportion of read libraries comprised of plasmids, phage, and genomic DNA (Fig. 1). Honey bee metagenomes were significantly enriched in plasmids compared to either commercial (TukeyHSD p < 0.01) or wild bumble bees (TukeyHSD p < 0.01). These findings indicate that the bacterial communities associated with honey bees harbor a larger plasmid load compared to bumble bees. This is consistent with previous research by Robinson et al.^23^, which found that plasmids, including those encoding AMR genes, to be widespread in honey bees. Given the longstanding practice of managing honey bees with antibiotics, the enrichment of plasmids in honey bee microbiomes may reflect a selective pressure favoring plasmids, especially those encoding AMR. Alternatively, this difference could be a result of differences in the overall abundance and diversity of gut bacteria associated with bumble and honey bees^10,24,25^.

**Figure 1.**
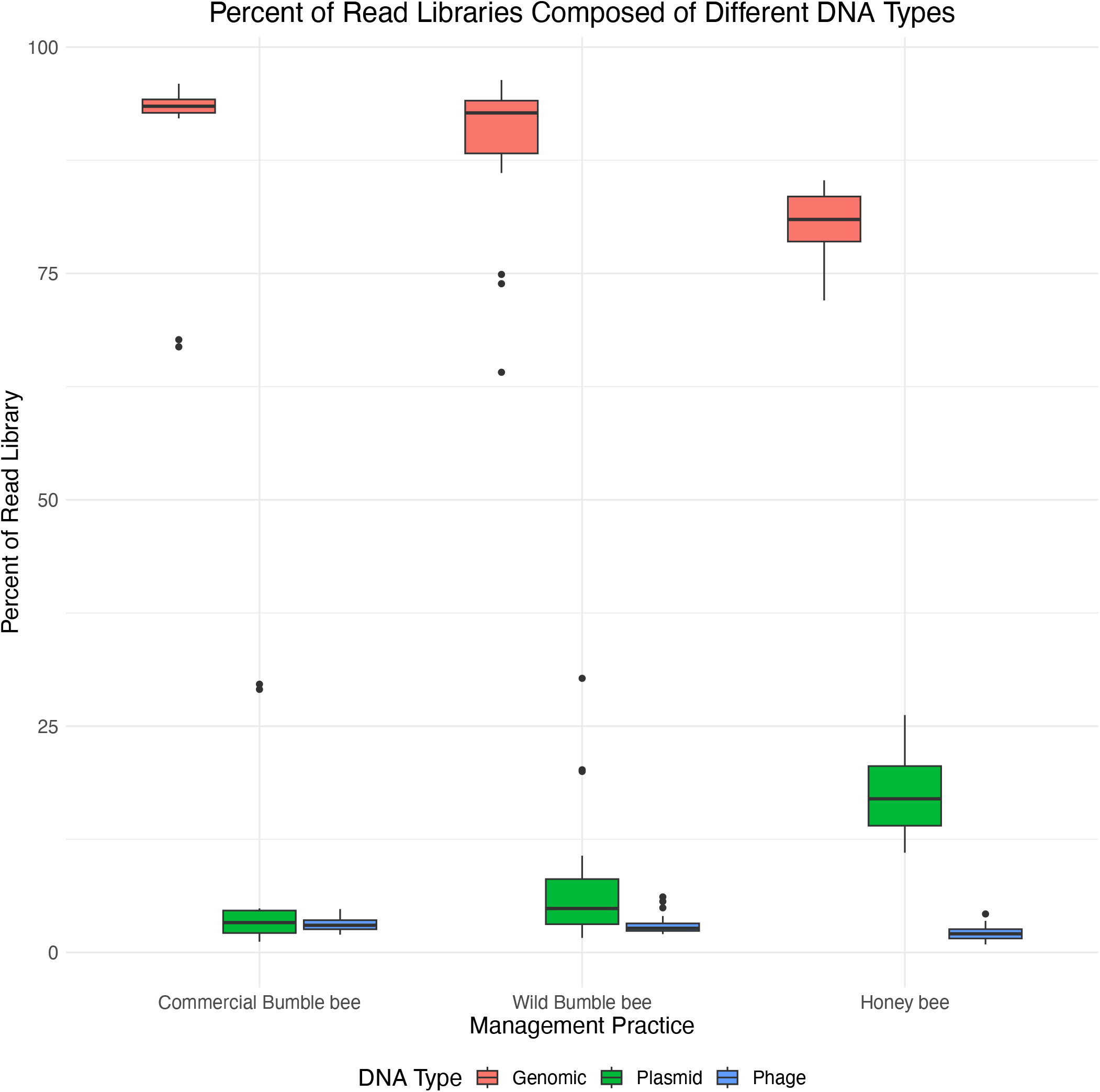
Figure describing proportion of metagenomic read libraries composed of genomic (red), plasmid (green), or phage (blue) DNA. Compared to either bumble bee population, more of the reads found in honey bee metagenomes are predicted to come from plasmids.

### Honey bee Plasmids Encode a High Diversity of AMR Genes

Next, we wanted to examine the diversity of AMR genes across bee populations and DNA types (Fig. 2). Honey bees exhibited the greatest number of unique AMR genes (Wild vs Honey TukeyHSD p < 0.01; Commercial vs Honey TukeyHSD p < 0.01), followed by wild bumble bees (Wild vs Commercial TukeyHSD p < 0.05), with the majority of honey bee AMR genes located on plasmids (TukeyHSD p < 0.01). No AMR genes were identified in phage-encoded DNA. We interpret the elevated diversity of AMR genes in honey bee plasmids to align with results from Tian et al.^7^, who found that tetracycline resistance was prevalent in honey bee associated bacteria and proposed that long-term antibiotic exposure has selected for resistant honey bee gut microbiota. Further, our results suggest that plasmids may play a key role in maintaining and disseminating AMR genes in the honey bee microbiome.

**Figure 2.**
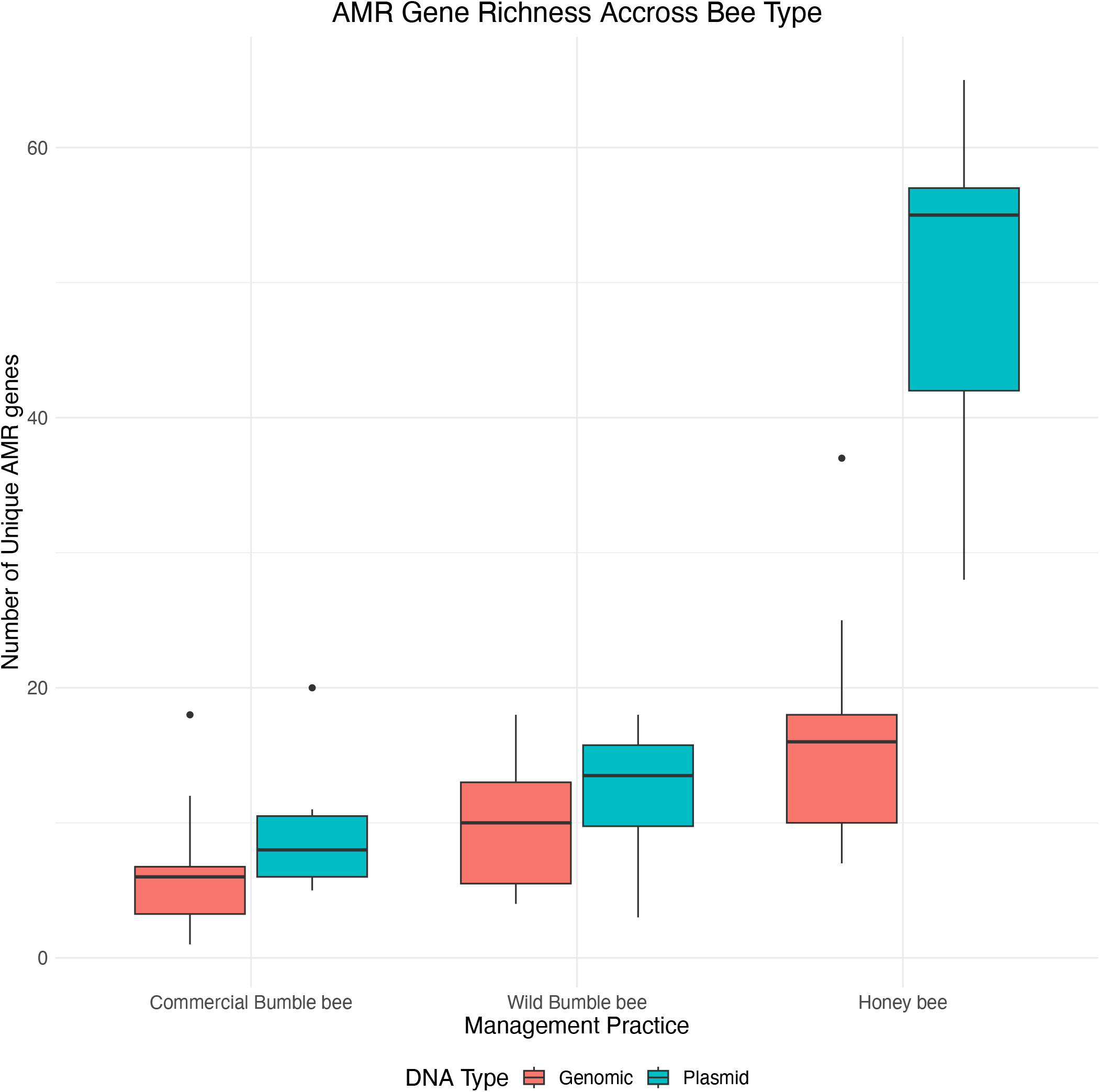
Boxplots illustrating how the number of unique antimicrobial resistance (AMR) genes differs in response to bee species and DNA type. Honey bee microbiomes house a greater diversity of AMR genes, compared to either bumble bee species. Most of the honey bee AMR genes predicted are encoded on plasmids.

### Abundance of AMR Genes Differs Modestly Between Bee Populations

Initially, we hypothesized that honey bees and managed bumble bees would harbor the highest (normalized) abundance of AMR genes due to routine antibiotic use. In contrast, we found only modest differences in overall AMR abundance between populations (Fig. 3). Commercial bumble bees had the lowest abundance of aminoglycoside (Commercial vs. Wild Bumble bee p < 0.01; Commercial vs. Honey bee p < 0.01) and macrolide-lincosamide-streptogramin (MLS) resistance genes (Commercial vs. Wild bumble bee p < 0.05; Commercial vs. Honey bee p < 0.05). No significant differences in abundance were observed between wild bumble bees and honey bees for either antibiotic class (aminoglycosides: p = 0.88; MLS: p = 0.82).

**Figure 3.**
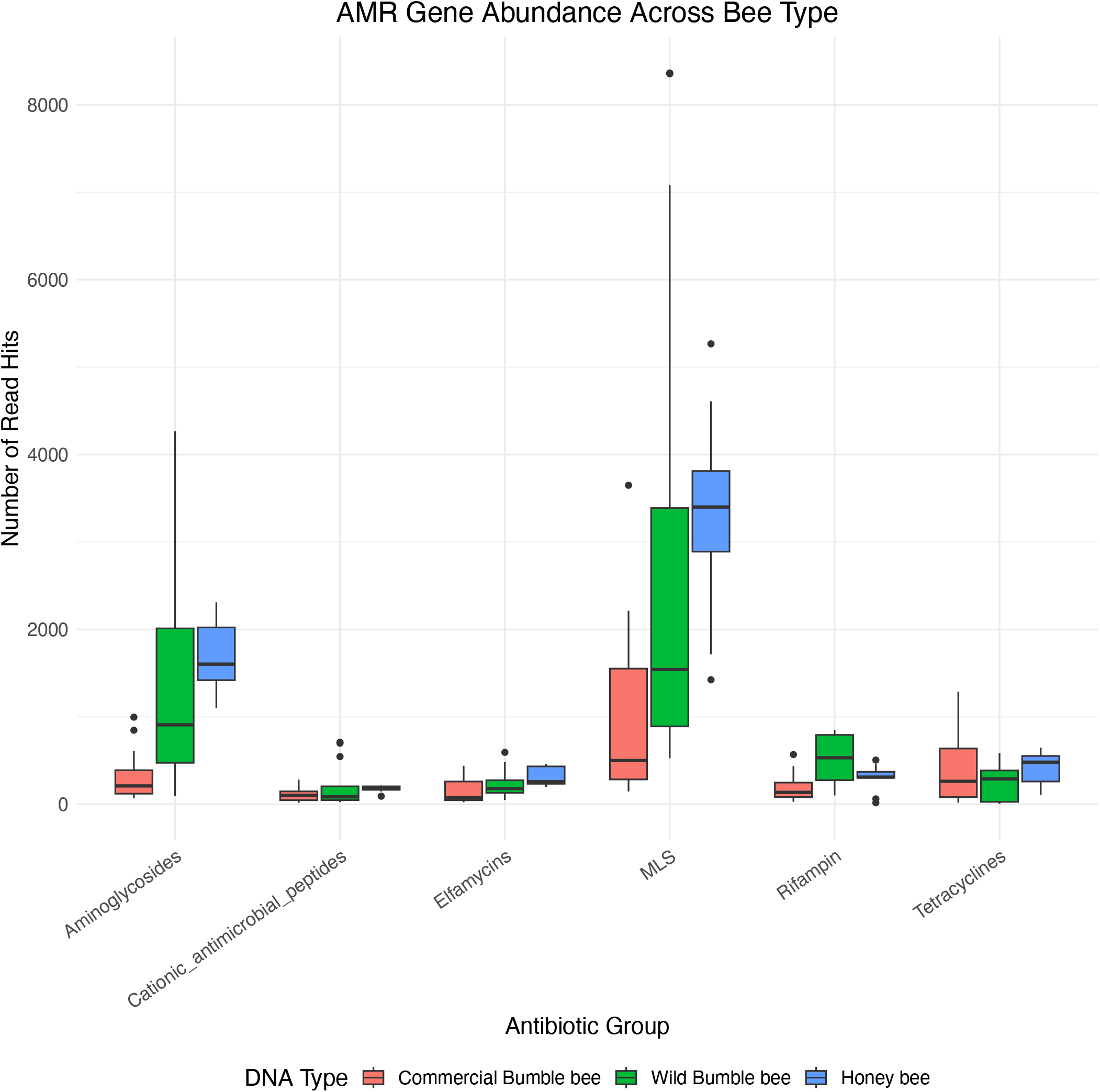
Figure describing how AMR abundance differs in respect to bee species. AMR genes are grouped according to the antimicrobial compounds they are predicted to confer resistance to (x-axis). The y-axis represents total number of normalized reads mapped. Overall, AMR genes are more abundant in honey bees and wild bumble bees, compare to commercial bumble bees, though these differences are typically modest.

While these results could suggest that antibiotic usage does not equate to higher abundance of resistance genes, it is important to view this result in a broader context. The honey bees sampled here were not actively being treated with antibiotics. As a result, there was likely no current pressure increasing the abundance of resistance genes. Alternatively, while there are only modest differences in the AMR abundance, there could still be important differences at the transcription level which are not captured here.

### Mechanisms of AMR Differ by Bee and DNA Type

Given the widespread use of tetracyclines in apiculture, we chose to further examine the mechanisms underlying tetracycline resistance across bee populations (Fig. 4). Despite similar total abundance of tetracycline resistance genes, the mechanisms and genomic locations of these genes differed substantially. Bumble bee-associated bacteria primarily encoded resistance via mutations in the 16S rRNA subunit, whereas honey bee-associated bacteria utilized plasmid-encoded efflux pumps and ribosome protection proteins. These mechanistic differences likely represent the legacy of previous antibiotic treatments and could have important impacts on how these bacterial communities respond to future antibiotic exposure. Future work could test this by measuring how these mechanistic differences change during active antibiotic exposure. Such an experiment could test if plasmid-encoded efflux pumps proliferate, and perhaps even jump hosts, following antibiotic treatment.

**Figure 4.**
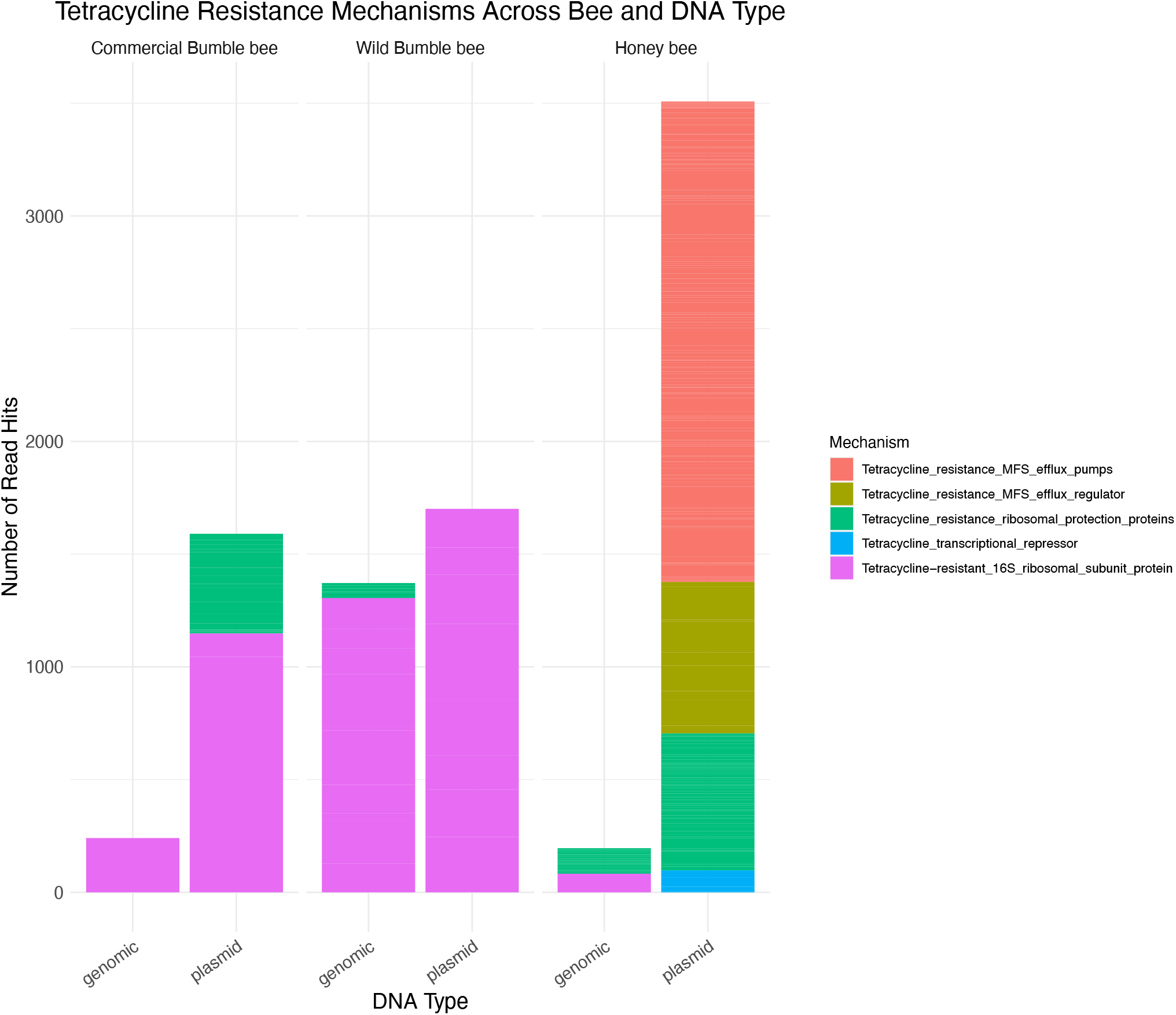
Stacked bar plots showing how the mechanisms of tetracycline resistance differ between bee and DNA type. While bumble bee bacteria appear to develop tetracycline resistance via altering the drug target (16S sequences), honey bees appear to leverage efflux pumps as a resistance mechanism. Moreover, those efflux pumps are found exclusively on plasmids.

## Conclusions

The emergence of antimicrobial resistance (AMR) in human health and agricultural systems is a global health concern. AMR surveillance through genomic methods can inform the ecological context of resistance genes in natural populations. In this work, we provide a glimpse into the diversity and abundance of AMR genes across bee species and populations under varied management. Our results show that the bacteria housed by managed honey bees encode a greater diversity of AMR genes, relative to either commercial or wild bumble bees, and that many of these resistance genes are likely present on plasmids. We also show that honey bee bacteria, which are frequently exposed to tetracycline antibiotics, employ wholly different mechanisms to resist those antibiotics compared to bumble bee bacteria. Overall, we interpret this data as showing that honey bees have a strong legacy effect of previous antibiotic treatment.

## Acknowledgements

We thank Dr. Hotchkiss, Dr. Poulain, and Dr. Forrest for their correspondence and willingness to share data. We also thank the Vannette lab for comments on the manuscript.

## Data Availability

All scripts associated with this manuscript and analysis are available on Github (https://github.com/dsbard/AMR_2025).

